# OCCURRENCE OF THE BRACHIOPOD *TICHOSINA* IN DEEP-SEA CORAL BOTTOMS OF THE CARIBBEAN SEA AND ITS PALEOENVIRONMENTAL IMPLICATIONS

**DOI:** 10.1101/2020.06.24.168658

**Authors:** Alexis Rojas, Adriana Gracia, Ivan Hernández-Ávila, Pedro Patarroyo, Michał Kowalewski

## Abstract

Despite its importance as the larger component of the modern and Cenozoic brachiopod faunas in the Caribbean region, the ecology and habitat preferences of the terebratulid *Tichosina* remain poorly understood. We compiled field observations from multiple sites in the Caribbean of Colombia (i.e., San Bernado Bank, Bahia Honda-Guajira, Puerto Escondido, Joint Regime Area Jamaica-Colombia) and data from the R/V Pillsbury program, indicating that *Tichosina* may have close ecological ties with deep-water corals. In addition, we reviewed literature sources on Cenozoic sediments in the Dominican Republic and found tentative evidence that such ecological ties could have existed since at least the Pliocene. These observations are reminiscent of the *Gryphus*-anthozoan association observed in the modern Mediterranean continental margin. Understanding to what extent the brachiopod *Tichosina* is linked to deep-water habitats has implications for the recognition of deep-water macrobenthic communities in the Cenozoic rock record of the Caribbean.

## INTRODUCTION

Deep-sea brachiopods have been recorder in the Caribbean Sea by several expeditions taking place in this region, i.e., the U.S. Coast Survey vessel Blake (late 1870s), U.S. Fish Commission Albatross (1880s), R/V Oregon and R/V Oregon II (1950’s to 1970s), R/V Gerda (1960s) and R/V Pillsbury campaigns (1960s and 70s) (see also Lutz and Ginsburg, 2007) provided abundant material to describe their taxonomy, general bathymetric and geographic distribution (Cooper, 1977; Logan, 1990). Micromorphic forms (< 10 mm in shell length), including *Argyrotheca* and *Terebratulina* species, are known to inhabit caves and other cryptic habitats (Jackson et al., 1971; Asgaard and Stentoft, 1984; Logan, 1975; Richardson, 1997) but under low-intensity pressures they can also occur on relatively exposed surfaces (Logan, 1977). In contrast, medium to large size brachiopods, comprising eighteen *Tichosina* species occurring in the Caribbean Sea and the Gulf of Mexico (Cooper, 1977; Rojas et al., 2015), have not been studied extensively and their taxonomy and ecology remain poorly understood (Harper et al., 1997; Oyen et al., 2000; Harper and Donovan, 2007; Emig et al., 2013). These brachiopods are known to occur in the depth range between 100 and 200 m (Cooper, 1977), most likely below the photic zone, but their potential association with deep-corals habitats, developed at depths greater than 70 m (Roberts, 2006; Ramirez-Llodra et al., 2010), has not been evaluated. In contrast, fossil brachiopods in the Caribbean region has been extensively studied with stratigraphic, taxonomic (Harper, 2002; Harper and Donovan, 2007; Harper and Pickerill, 2008), and biogeographic purposes (Shemm-Gregory et al., 2012; Sandy and Rojas, 2018, Rojas and Sandy, 2019).

Although deep-sea corals across the Caribbean Sea have been widely studied during the last decades (Cairns, 1979, 2007; Lutz and Ginsburg, 2007; Davies and Guinotte, 2011; Santodomingo et al., 2013; Hernandez-Avila, 2014), information on the brachiopod faunas associated with these coral-supported environments is limited (Cooper, 1979). These light-independent structures, including banks, bioherms, lithoherms and other coral-supported structures generated by azooxanthellate corals (i.e. corals lacking symbiotic dinoflagellates), provide protection and food, as well as breeding, spawning, resting and nursery areas for several species of invertebrates and vertebrates (Roberts, 2006; Tursi et al., 2004). For instance, a deep-water coral bank located in the Southwestern Caribbean of Colombia is known to provided habitat for at least 118 species of fishes and invertebrates, including echinoderms, crustaceans, gastropods, bivalves, bryozoans and brachiopods (Reyes et al., 2005; Rojas et al., 2015). However, the distinction of these deep-water macrobenthic communities in the Cenozoic rock record is challenging (Squires, 1964; Mullins et al., 1980; Kano et al., 2007) and water-depth interpretations of some biosediments in the region remains controversial (McNeill et al., 2008).

In this paper, we compiled field observations on brachiopods from multiple deep-shelf settings in the Caribbean of Colombia and available data from the R/V Pillsbury sampling program across the Caribbean (1966 to 1972), indicating that *Tichosina* may have ecological ties with deep-water coral habitats since it has been recorded consistently in trawl samples containing structure forming and suspected habitat-forming corals. In addition, we discussed literature sources providing tentative evidence that such brachiopod-anthozoan ties could have existed at least since the Pliocene. Understanding to what extent the brachiopod *Tichosina* is linked to deep-coral habitats through the modern Caribbean has implications for both Tertiary geology and paleoecology in this region.

## MATERIAL AND METHODS

### DATA

Three main data sources have been used in our study: literature sources, field collection observations, and databases. First, we examined uncatalogued *Tichosina* shells and fragments deposited at the Museo de Historia Natural Marina of the Instituto de Investigaciones Marinas y Costeras (INVEMAR, Colombia). The materials were collected during the MACROFAUNA II (2000), MARCORAL (off San Bernardo Archipelago), ANH-I (Bahia Honda-Guajira, off Cartagena and Puerto Escondido) and ANH-Jamaica (northwestern Macondo Guyot) projects that were focused on characterizing deep-water coral and soft-bottom communities and associated populations in the continental shelf of Colombia. These field surveys resulted in the recognition of a deep-water coral structure called San Bernardo Bank (Lutz and Ginsburg, 2007), as well as others coral-dominated habitats (see Reyes et al., 2005, Santodomingo et al., 2007, Santodomingo et al., 2013) in the Caribbean continental shelf of Colombia. Then, we used a dataset on the benthic macrofauna from the R/V Pillsbury sampling program (1966 to 1972), including corals, mollusks and echinoderms, assembled from databases of the Natural Museum of Natural History, Smithsonian Institution and the Marine Invertebrate Museum, Rosenstiel School of Marine and Atmospheric Science (MIN-RSMAS) (Hernández-Ávila, 2014). Brachiopod data from the R/V Pillsbury sampling program were gathered from an extensive monograph on modern Caribbean brachiopods elaborated by Cooper (1977). Finally, we partially inspected the Cooper’s collection (Cooper, 1977) deposited at the Smithsonian National Museum of Natural History for direct evidence of *Tichosina* shells growing on deep-water corals.

### METHODS

Brachiopod shells and fragments from the MACROFAUNA II, MARCORAL, ANH-I and ANH-Jamaica projects were handpicked and examined under a binocular microscope. Complete specimens of *Tichosina plicata* were counted and measured. As defined in Rojas and Sandy (2019), a complete specimen includes any shell sufficiently complete to measure its full dimensions (i.e. length and width). A taxonomic description of the brachiopods studied here is provided in Rojas et al. (2015). Because the San Bernardo Bank is the most conspicuous deep-water coral structure in the Southwestern Caribbean of Colombia (Lutz & Ginsburg 2007), we used the total samples from the MARCORAL cruise, carried out between April 22 and May 2, 2005 on board R/V Ancon off the San Bernardo Archipelago (Reyes et al., 2005), to illustrate variations in sampling effort, total number of brachiopod specimens and sediment grainsize at each station (Fig. 1). The sampling methodology of this survey was previously described by Rangel-Buitrago & Idárraga-García (2010).

**Figure 1.**
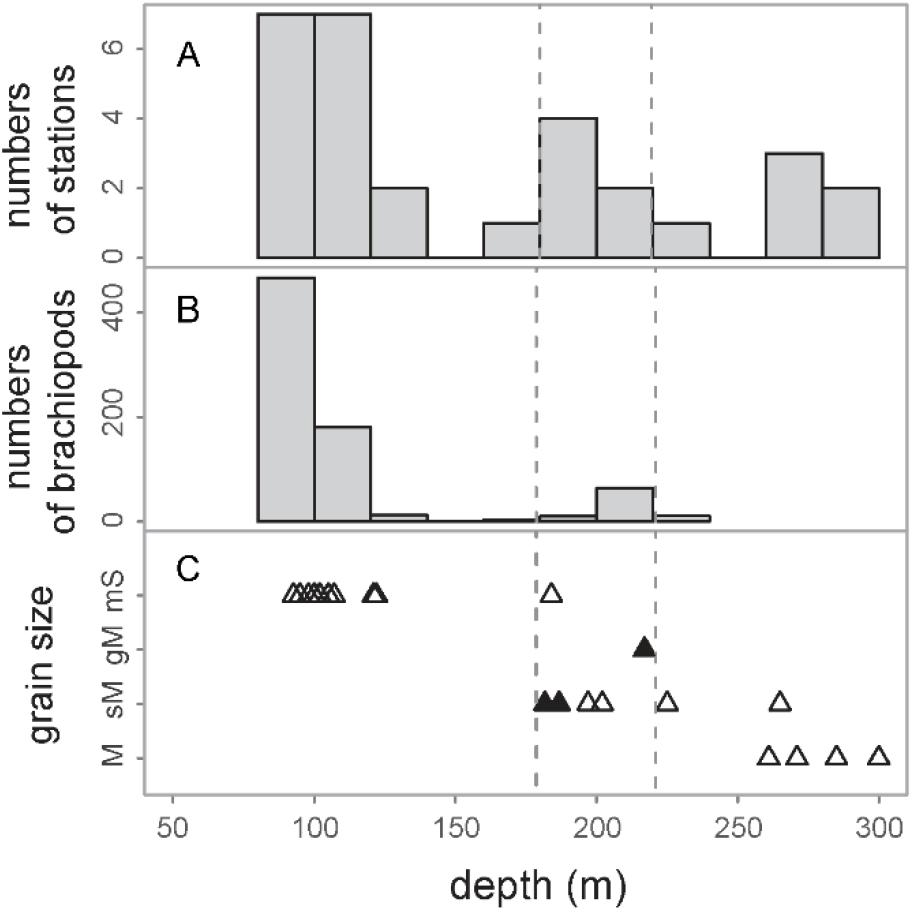
Sampling data from the INVEMAR Marcoral project carried out offshore San Bernardo, Colombia. A, Total number of stations; B, Total number of brachiopods, including *Terebratulina cailleti* (503 specimens), *Argyrotheca barrettiana* (208 specimens) and *Tichosina plicata* (6 specimens); C, Sediment grain size per station. Stations were *T. plicata* was recovered are indicated in black. Abbreviations for texture classes are: mud (M), sandy mud (sM), muddy sand (mS), sand (S) and gravelly mud (gM). Depth distribution of the San Bernardo Bank indicated by a segmented line.

The assembled dataset on the benthic macrofauna from the R/V Pillsbury sampling program was used to reconstruct the habitat of *Tichosina* spp. across the Caribbean Sea. The brachiopod specimens from these expeditions were described in the Cooper’s monograph (Cooper, 1977) and are housed at the Paleobiology Collections of the National Museum of Natural History (NMNH). Cooper’s monograph was focused on brachiopod taxonomy and it does not provide information on their interactions. Although all the materials recovered during the Pillsbury expeditions were divided into major taxonomic groups and placed at specialized collections, AR inspected some available brachiopods in the wet brachiopod collections housed at the Smithsonian National Museum of Natural History Home (NMNH) for any remaining evidence on the presumed *Tichosina*-anthozoan relationship.

The nonmetric multidimensional scaling (Kruskal, 1964) was used to explore distribution patterns of *Tichosina*, structure-forming and potential habitat-forming corals across the R/V Pillsbury sampling stations. It was performed using census data at the genus taxonomic-level and the Bray Curtis distance measure after removing rare taxa that provide limited interpretative value (Poos and Jackson 2012) on the occurrence of *Tichosina*. This ordination analysis was performed using the R-package *vegan* version 2.5-1 (Oksanen et al., 2018). In addition, we created a bipartite network representation of the census data gathered from R/V Pillsbury sampling stations (Data S1) and used a clustering analysis called Infomap (Rosvall and Bergstrom, 2008; Calatayud et al. 2019) to identify the potential impact of removing rare taxa in the multivariate analysis and further explore the modular structure of the benthic macrofauna. This community detection approach allows us to identify clusters of highly connected stations and taxa. We performed the network analysis using the MapEquation framework, available online at http://www.mapequation.org.

## RESULTS

### *TICHOSINA* IN DEEP-SEA CORAL BOTTOMS OF THE COLOMBIAN CARIBBEAN SEA

The combined *Tichosina* material from the INVEMAR projects comprises live and dead specimens, including 55 conjoined shells and 24 disarticulated valves. This shell material represents 10 sampling stations from 5 different sites along the Caribbean continental shelf of Colombia. These sites are located offshore Guajira Province, Bolivar Province, Córdoba Province, and the northwestern Macondo Guyot in the Joint Regime Area (JRA) Jamaica-Colombia (Gutiérrez-Salcedo et al., 2015) (Table 1) (Fig. 1). Deep-sea *Tichosina* was recorded in the depth range between 120 and 500 m and all studied localities register one single species, *T. plicata* (Rojas et al., 2015). Well-preserved dead specimens were found in sandy and gravelly litho-bioclastic mud (Rangel-Buitrago and Idárraga-García, 2010) (Fig.2). Although the dominant coral species in the San Bernardo Bank is *Madracis myriaster* (Reyes et al., 2005; Santodomingo et al., 2007), a few live specimens were found attached to dead and live branching corals *Madracis asperula, Madracis* sp., also in fragments of the corals *Agaricia* sp. and *Anomocora fecunda* (Fig. 2). Dead specimens of *Tichosina* were observed attached to the stony coral *Caryophyllia* sp. Offshore San Bernardo Archipelago, the deep-sea brachiopod fauna includes also three micromorphic forms, *Terebratulina cailleti* (503 specimens)*, Argyrotheca barrettiana* (208 specimens), and *Terebratulina latifrons* (1 specimen) (Rojas *et al.* 2015). These brachiopods were common on the continental shelf with only a few occurring on the shelf break and associated to the deep-water coral structure.

**Figure 2.**
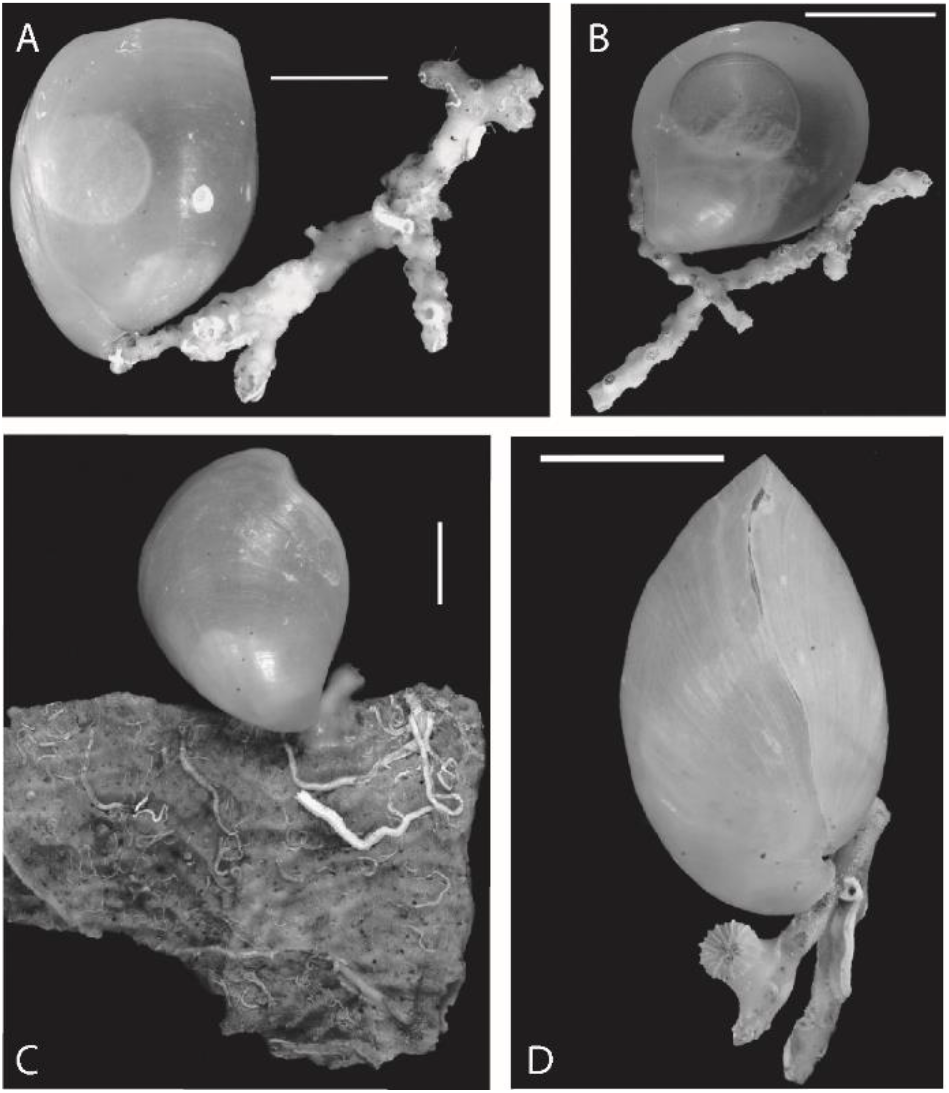
Specimens of *Tichosina plicata* attached to deep-water corals from the San Bernardo Bank. A, *Madracis asperula*; B, *Madracis* sp.; C, Dead *Agaricia* sp. and recruit of Caryophylliidae; D, *Madracis* sp. and recruit of Caryophylliidae. Scale bars 10 mm.

Dead and living specimens of *Tichosina* sp. in environments dominated by solitary scleractinians were also observed in Bahia Honda-Guajira (265 m ANH I Project, Sts. E253 and E254), off Cartagena (500 m ANH I Project, St. E264), off Puerto Escondido (500 m ANH I Project, St. E267) and the northwestern Macondo Guyot (320 m ANH-Jamaica, St. E296). Geological and biological evidence suggest the presence of deep-coral structures in all those sites.

### *TICHOSINA* AND THE BENTHIC MEGAFAUNA FROM THE R/V PILLSBURY STATIONS

A total of five species of *Tichosina* were reported from 28 stations of the R/V Pillsbury sampling program, including *T*. *plicata* (=*T. dubia*), *T. obesa, T. truncata, T. bartletti* and *T. cubensis.* These five species represent 28 % of all species assigned to this genus that have been reported to date from the Caribbean and the Gulf of Mexico (Cooper, 1977). Because most of the species of this genus appear to be synonymous (Emig et al., 2013; Rojas et al., 2015), our analises are performed at the genus level. Occurrences of the brachiopod *Tichosina* in the R/V Pillsbury data is limited to only 28 out of 318 stations with coral records. The scarcity of *Tichosina* may likely reflect the sampling method (e.g. otter trawls), which was not optimal for capturing small and fragile specimens. Remarkably, all sampling stations from the R/V Pillsbury sampling program recording *Tichosina* brachiopods, as reported in Cooper (1977), also register structure-forming and potential habitat-forming corals in our compilation. The megafauna associations at those stations represent relatively diverse deep-water benthic communities, including corals (102 spp), gastropods (63 spp), echinoids (40 spp), asteroids (37 spp) and crinoids (15 spp) (Table 2). The diverse material collected from those stations includes the structure-forming coral *Madrepora oculata*, as well as a number of potential habitat-forming corals (*sensu* Lutz and Ginsburg, 2007), including *Acanella* sp., *Diodogorgia* sp., *Riisea paniculata*, *Swiftia exserta*, *Parantipathes* sp., *Coenosmilia arbuscula*, *Madracis asperula*, *M. formosa*, *Nicella americana*, *N. guadalupensis*, *Neospongode*s sp., *Oxysmilia rotundifolia*, *Thalamophyllia riisei*, *Eguchipsammia cornucopia,* and *Polymyces fragilis*. Each of those coral species occurs in 4 to 11 % of the total stations.

The ordination diagram resulted from the non-metric multidimensional scaling analysis (Fig. 3A) does not show a clear distinction between habitat-forming corals and other anthozoans across the R/V Pillsbury sampling stations. However, *Tichosina* plots into the area of higher dot density, that comprises both habitat-forming (i.e., *Madracis*) and potential habitat-forming species (i.e., *Swiftia, Riisea* and *Nicella*). The well-known structure-forming coral *Madrepora* plots outside this region. Estimation of the Besag’s *L*-function (Besag, 1977) indicates significant clustering for the dot pattern resulted from the ordination analysis. Network clustering analyses based on the reduced dataset used in the multivariate analysis as well as the complete dataset that includes rare taxa show a similar modular structure, i.e., despite removing rare taxa, we were able to retrieve similar modules or communities (Fig. 3B). The reference solution, obtained by clustering the assembled network representing the R/V Pillsbury campaign, includes 29 modules that comprise both sampling stations and genera (Data S2). In this network partition, *Tichosina* is clustered with deep-water brachiopods as well as solitary, attached or free-living corals, including the potential habitat-forming *Parantipathes* and *Polymyces* (*sensu* Lutz and Ginsburg, 2007). The three larger modules delineating in the structure of this benthic megafauna are showed in figure 4. Finally, the partial inspection of the Cooper’s collection (Cooper, 1977) deposited at the Smithsonian National Museum of Natural History resulted in a single wet sample from Venezuela (uncatalogued, Pillsbury sampling station P736) of *Tichosina* growing on an undetermined coral fragment (Fig. 4).

**Figure 3A.**
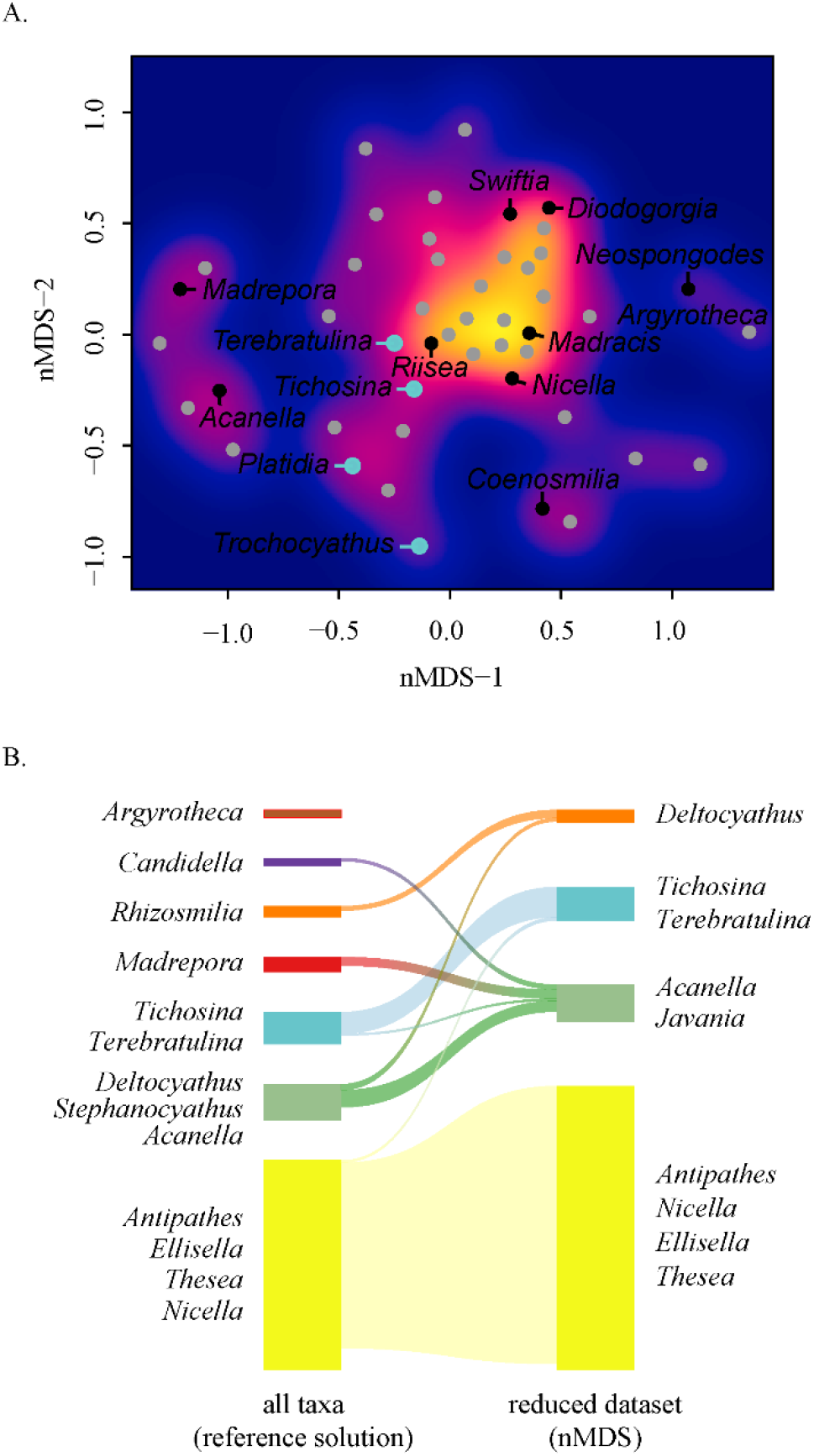
Non-metric multidimensional scaling plot for combined coral and brachiopod genus census data from the R/V Pillsbury campaigns. Only common genus (recorded in 3 or more samples) and samples in which 3 or more species are recorded were used in this ordination. The stress value for the three-dimensional analysis is 0.13, and the *R*-squared value for the linear fit of observed dissimilarity and ordination distance is 0.9. **3B**— Alluvial diagram showing the differences between the two partitions derived from network representations of the total census data from R/V Pillsbury sampling stations and the reduced dataset generated after removing rare taxa.

## DISCUSSION

### *TICHOSINA* IN DEEP-SEA CORAL BOTTOMS OF THE COLOMBIAN CARIBBEAN SEA

The Macrofauna II and Marcoral sampling programs targeted deep-sea corals structures and the sampling methodologies were not suitable for recovering fragile material like the *Tichosina* shells. Consequently, the available material is represented largely by shell fragments and its abundance, based on complete specimens, is likely to be severely underestimated. In contrast, the relatively strong shells of the micromorphic brachiopods in our samples, *Argyrotheca and Terebratulina*, were dominated by conjoined specimens. This negative effect of some sampling methodologies on selected taxa has been observed in modern ecological studies (Costello et al., 2017). Remarkably, complete specimens and abundant fragments of *T. plicata* were yet recovered at sites where branching *Madracis* spp. and other potential habit-forming corals were present. The azooxanthellate *Madracis* corals is considered the main framework builder of deep-water coral structures in the Colombian Caribbean (Santodomingo et al., 2007). Other scleractinian corals considered to represent habitat-forming species and a potential habitat to *Tichosina* includes *Anomocora fecunda*, *Coenosmillia arbuscula*, *Eguchipsammia cornucopia*, *Thalamophyllia riisei,* and *Javania cailleti* (Reyes et al., 2005; Santodomingo et al., 2007). The consistent cooccurrence of live specimens of both *Tichosina* and deep-water corals points to a potential *Tichosina*-anthozoan associations in the Caribbean continental shelf of Colombia. Although lack of *in situ* observations prevent a detailed description of such an interaction, it is likely that *Tichosina* occurs on the periphery of the coral structures, i.e., off-mound (Rosso et al., 2010). Well-preserved specimens of *T. rotundovata* has been also reported in association with deep-water coral structures in the little Bahama Bank at 485-496 m water depth (Cooper, 1977, p. 21). To evaluate at what extent are the local occurrences of *Tichosina* controlled by the presence of deep-water coral habitats across the modern Caribbean we examined benthic megafauna of the R/V Pillsbury survey. Because most of the species of *Tichosina* appear to be synonymous (Emig et al., 2013; Rojas et al., 2015), our analises are performed at the genus level.

### TICHOSINA AND THE BENTHIC MEGAFAUNA OF THE R/V PILLSBURY SAMPLING PROGRAM

Our data compilation suggests that *Tichosina* spp. are associated to deep-water benthic communities dominated by corals and other megafaunal elements characteristics including gastropods, echinoids, asteroids and crinoids. The coral material reported in those sampling stations includes structure-forming and potential habitat-forming corals. Those corals appear to provide the optimum structural habitat for *Tichosina* species to grow across the Caribbean. The number of azooxanthellate corals in the Caribbean region is higher on the continental slope (Hernández-Ávila, 2014), which coincides with the depth distribution of *Tichosina* spp. (Cooper, 1977; Harper et al., 1995; Harper, 2002). The commensal relationship between *T. floridensis* and benthic foraminifera occurring at depths of 120-180 m on the west Florida shelf illustrates the relationship of these brachiopods with the deep-water benthos (Zumwalt and Delaca 1980). Although the original material recovered from the R/V Pillsbury survey has been divided into major zoological groups and housed at specialized collections, we inspected some available wet samples of brachiopods at the National Museum of Natural History (NMNH) and found remains at station P736 from Venezuela that are reminiscent of the substrate recovered at the San Bernardo Bank. Despite that all samples from the R/V Pillsbury survey containing *Tichosina* brachiopods also includes known or potential habitat-forming corals, results from the NMS indicates there is not a preferred coral species for *Tichosina* to growth. Different deep-coral species appears to offer a suitable environment for *Tichosina* to growth across the Caribbean (Fig. 4). However, networks clustering analysis points to both *Parantipathes* and *Polymyces* as the most suitable corals for *Tichosina* to growth. This finding should be further corroborated with field observations.

**Figure 4.**
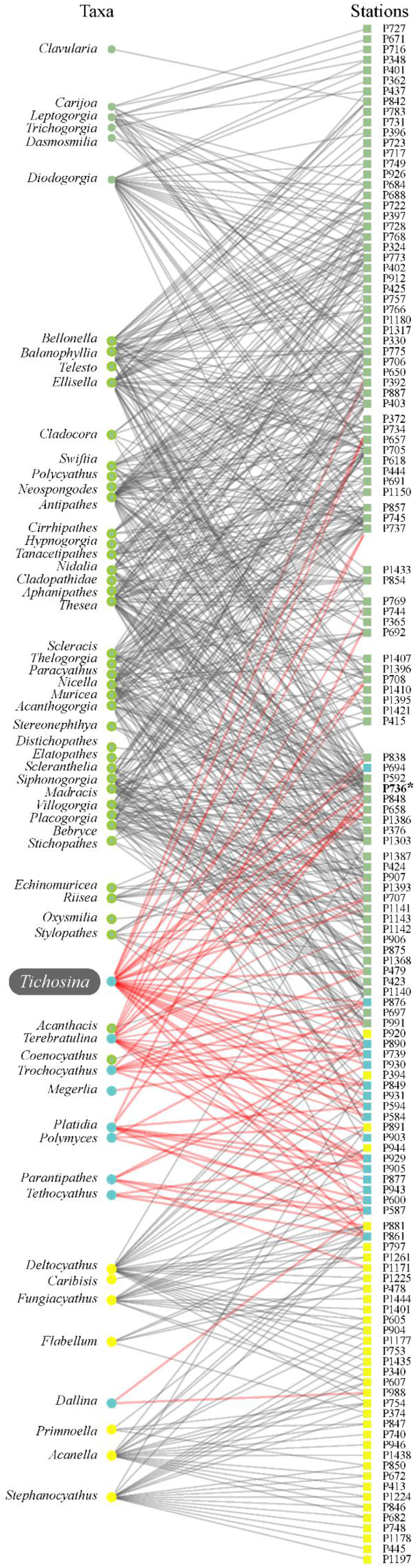
Bipartite representation of the census data from R/V Pillsbury sampling stations with nodes, species and stations, colored by their module affiliation. (*) Sample with *Tichosina* shell growing on undetermined coral fragment in the Copper’s collection. Only the three largest modules are illustrated.

Coral-brachiopod associations in deep-water environments are known from temperate zones, where deep-coral banks provide widespread habitats for abundant and diverse benthic organisms (Teichert, 1958; Squires, 1964; Logan, 1979; Rosso et al., 2010). Similarly, based on direct observations and dredging, Emig (1987, 1989) reported that Mediterranean continental-margin environments (depths of 110-250 meters) with strong bottom currents, low sedimentation rates and high nutrient delivery harbor higher density populations of *Gryphus vitreus* (Emig, 1997). The San Bernardo Bank is similar to the Mediterranean settings in that it is influenced by an undercurrent called the Panama-Colombia Countercurrent (Andrade et al., 2003). The local nutrient enrichment potentially associated with this current is unknown. Moreover, *Gryphus* and *Tichosina*, represent Cenozoic genera that are similar in shape and size (Harper and Pickerill, 2008). *Gryphus* is well represented in the Eocene of Cuba but is absent in the modern Caribbean Sea (Cooper, 1979). A recent study of modern surface sediments in a deep-water coral mound located at Santa Maria di Leuca (Mediterranean) confirmed the association of *G. vitreus* with coral-supported structures (Rosso et al., 2010). These previous studies and the findings reported here suggest consistently that populations of *Tichosina* are likely to concentrate along the continental margins of the Caribbean, where suitable habitats for deep-water corals exist (Hernandez-Avila, 2014).

Understanding to what extent *Tichosina* spp. are linked to deep-coral habitats through the modern Caribbean has several implications for both Tertiary geology and paleoecology in the region. We re-examine the occurrence of *Tichosina* with deep-water corals in the Neogene Mao Adentro Limestone of the Dominican Republic. Even though this geological unit is well-known in terms of stratigraphy and paleontological content, controversy remains over whether some of its carbonates are shallow- or deep-water in origin (McNeill et al., 2008), with obvious implications for our understanding of the regional sea-level history in the region.

### *TICHOSINA* FROM DEEP-WATER CORAL HABITATS IN NEOGENE DEPOSITS IN THE DOMINICAN REPUBLIC

In a detailed stratigraphic survey, Saunders et al., (1986) observed several sedimentological (e.g. channeling, load features, flame structures and silt casts) and biological (e.g. deep-water assemblages dominated by pteropod mollusks and deep-water corals) features indicating a deeper-water environment (>100 m in depth) for the base of Mao Formation at the Rio Gurabo section. A taxonomic study of the corals carried out by Cairns and Wells (1987), based on 1590 specimens representing 20 species, suggested an even deeper depositional environment (>200 m in depth) for certain parts of the Mao Formation. Those potentially deep-water environments include localities NMB 15823 and 15827-15029 from stratigraphic horizons between 650 and 750 m in the Rio Gurabo section (Cairns and Wells, 1987, table 1; Saunders et al., 1986, fig. 4). A deep-water origin for those rocks is supported by the occurrence of *Pourtalocyathus hispidus* (Pourtalès 1878), a deep-water coral species in the modern Caribbean (Cairns and Wells, 1987), and the composition of the mollusk assemblages (Saunders et al., 1982).

Terebratulid brachiopods mentioned briefly by Saunders et al. (1986) were later identified by Logan (1987) as *Tichosina*? *lecta*, a species described by Cooper (1979) based on Eocene material from Trinidad. The foramen shape and size of *Tichosina*? *lecta* is similar to those reported for other species of *Tichosina*, including the Eocene *Tichosina*? *trinitatensis* and a number of modern species including *T. pillsburyae*. *T. obesa*, *T. labiate*, *T. expansa,* and *Tichosina?bartletti* (Cooper, 1977, 1979). The uniplicate anterior margin of *Tichosina*? *lecta* is also a common feature in several modern species of *Tichosina* (Rojas et al., 2015). Logan (1987) also described *Tichosina* fragments gathered at the upper part of the Mao Adentro Limestone Member on the Rio Cana section, locality NMB 16884 (Saunders et al., 1987; Schultz and Budd 2008). This locality contained a diverse coral assemblage including genera recorded in modern deep-water habitats of the Caribbean (e.g. *Madracis* and *Manicina*). Corals species common in modern shallow waters were also present in the upper part of the Mao Adentro Limestone member in the Rio Cana section (Budd and Klaus, 2008). However, only a few of those coral remains appeared to have been preserved in life position (Saunders et al., 1982). Complete specimens of *Tichosina*? *lecta* showing post-mortem distortion and fracturing were also reported at the very bottom of the Mao Formation in the Rio Gurabo section, locality TU 1300 (Saunders et al., 1987; Logan, 1987). Saunders et al. (1986) identified an increase in the richness of planktonic foraminifera from the upper Gurabo Formation through the lower Mao Formation in the Rio Gurabo section. The latter include the stratigraphic horizons with the *Tichosina* remains mentioned above.

McNeill et al. (2008) re-interpreted the Mao Formation as a progressive shallowing sequence based on a regional interpretation of the Cibao Basin and the Bahamas, with both regions experiencing shallowing during the late Pliocene. The water-depth interpretation of the Mao Formation by McNeill et al. (2008) was established on the Rio Cana section, which is located more than 15 km west of the Rio Gurabo section. The Mao Formation at the Rio Gurabo section differs in terms of thickness of sediments, proportion of different facies, and paleontological records when compared to the Rio Cana section. The Mao Adentro Limestone *sensu* Saunders et al. (1982) in the Rio Cana section comprises ~350 m of thick bedded coral limestone and massive limestones with the bivalves *Ostrea, Lithophaga,* and shallow-water reef forming corals. In contrast, this member has not been identified satisfactorily in the Rio Gurabo section (see Saunders et al., 1982, fig. 3, Saunders et al., 1986, fig. 4). The potential sediments corresponding to this member in the Rio Gurabo section are ~100 m thick sequence of intercalated limestones and siltstones obscured by covered outcrop intervals and faults. As indicated above, those deposits are characterized by deep-water taxa including *Limopsis* sp., and the corals *Pourtalocyathus hispidus, Dendrophyllia cornucopia* and *Trochocyathus duncani* (Saunders et al., 1982, fig. 2; Cairns and Wells, 1987). Although the stony cup coral *Dendrophyllia* may not be a member of the deep-coral communities because its bathymetric distribution is generally lower (Emig personal communication, 2018), The genus *Trochocyathus* was clustered together with *Tichosina* in the network analysis of the modern Caribbean data (Fig. 4).

Based on these observations, the sediments from the Rio Gurabo section bearing *Tichosina* remains were likely to be deposited in deep-water settings (>100 m) below the photic zone. Our re-evaluation of the paleontological data of the Mao formation suggests that *Tichosina* may has been associated with deep-water coral communities since at least the Pliocene. Large terebratulids of the genus *Tichosina* have been reported in post-Paleocene sediments across the Caribbean basin (see Harper, 2002). Typically, the setting of those records is interpreted as a fore reef, an extension of the shallow-water coral environments (Harper et al., 1995; Harper et al., 1997; Harper et al., 2002). However, depth estimations for those deposits frequently exceed the deepest limit of the bathymetric distribution of modern zooxanthellate corals (Englebert et al., 2014). Fossil assemblages with *Tichosina* spp., as those described recently by Donovan et al. (2015), may represent depth-water benthic communities developed below the photic zone.

## CONCLUSIONS

Our observations from multiple sites on the Caribbean continental shelf of Colombian suggest that *T. plicata* inhabit deep-water coral habitats dominated by known and potential habit-forming corals (i.e., *Madracis* spp., *Anomocora fecunda*). Furthermore, compiled data from the R/V Pillsbury sampling program indicates that *Tichosina* spp. may have close ecological ties with deep-water corals since they were recovered consistently across trawl samples containing structure-forming (i.e., *Madrepora oculata*) and potential habitat-forming corals (e.g., *Acanella* sp., *Diodogorgia* sp., *Riisea paniculata*, *Swiftia exserta*, *Parantipathes* sp., *Coenosmilia arbuscula*, *Madracis asperula*, *M. formosa*, *Nicella americana*, *N. guadalupensis*, *Neospongode*s sp., *Oxysmilia rotundifolia*, *Thalamophyllia riisei*, *Eguchipsammia cornucopia,* and *Polymyces fragilis*). Furthermore, networks analysis points to *Parantipathes* and *Polymyces* as important habitat-forming corals for *Tichosina* to growth. Deep-water benthic communities dominated by solitary, attached and free-living corals appear to provide the optimum structural habitat for *Tichosina* spp. to growth across the continental shelves of the modern Caribbean Sea. A re-evaluation of the paleontological data from carbonates of the Mao Adentro Formation (Rio Gurabo section) in the Dominican Republic, suggests that *Tichosina* brachiopods may have been tied to deep-water corals habitats since at least the Pliocene (~3 Ma). Understanding to what extent the brachiopod *Tichosina* is linked to deep-coral habitats through the modern Caribbean Sea has several implications for both Tertiary geology and paleoecology in the region. Although the distinction of deep-water communities in the Cenozoic rock record is challenging, this potential brachiopod-Anthozoan association could be useful to distinguish those communities in the Cenozoic rock record and thus it would enhance our understanding of the sea-level history in the Caribbean basin.

## ACKNOWLEDGMENTS

We thank Nadia Santodomingo (Natural History Museum, London) for identification of corals and Miguel Martelo and Erika Montoya for support at the Marine Natural History Museum of Colombia. AR thanks Roger Portell (Florida Museum of Natural History) for advice and support. We thank Gustav Paulay (Florida Museum of Natural History), Liang Mao (Department of Geography, University of Florida) and Christian C. Emig (BrachNet) for their comments on earlier drafts of this report. AR was supported by a Jon L. and Beverly A. Thompson Endowment Fund. MARCORAL project was funded by Invemar -Colciencias (Project Code 2115-09-16649). Coral database from the Pillsbury expedition were provided with the permission of the Natural Museum of Natural History (Smithsonian Institution), the Marine Invertebrate Museum, and the Rosenstiel School of Marine and Atmospheric Science (MIN-RSMAS). IHA thanks Nancy Voss for providing the MIM-RSMAS data. IHA is supported by a Fundayacucho PhD grant E-223-85-2012-2, and Campus France grant 796045K.

## TABLE CAPTIONS

Table 1—Information on sampling stations were *Tichosina* spp. have been recovered in the Caribbean Sea by the INVEMAR projects and the R/V Pillsbury campaigns. Brachiopod data of the R/V Pillsbury campaigns from Copper (1977).

Table 2—Major components of the benthic megafauna from R/V Pillsbury stations were *Tichosina* spp. have been recovered in the Caribbean Sea.

## SUPPLEMENTARY MATERIAL

Data S1—Results of the clustering analysis on the network assembled in this study (R/V Pillsbury sampling stations). This standard file contains the reference solution. Each row begins with the module assignments of a node in a colon-separated format. Nodes within each module are sorted by the total amount of flow they contain. The decimal number is the amount of flow in each node. The clustering procedure was implemented using the Infomap algorithm (Rosvall & Bergstrom 2008).

Data S2—Results of the clustering analysis on the network assembled in this study (R/V Pillsbury sampling stations). This standard file contains the reference solution. Each row begins with the module assignments of a node in a colon-separated format. Nodes within each module are sorted by the total amount of flow they contain. The decimal number is the amount of flow in each node. The clustering procedure was implemented using the Infomap algorithm (Rosvall & Bergstrom 2008).

